# An optimized FM-index library for nucleotide and amino acid search

**DOI:** 10.1101/2021.01.12.426474

**Authors:** Tim Anderson, Travis J Wheeler

## Abstract

Pattern matching is a key step in a variety of biological sequence analysis pipelines. The FM-index is a compressed data structure for pattern matching, with search run time that is independent of the length of the database text. We present AvxWindowedFMindex (AWFM-index), an open-source, thread-parallel FM-index library written in C that is optimized for indexing nucleotide and amino acid sequences. AWFM-index is easy to incorporate into bioinformatics software and is able to perform exact match count and locate queries ~2-4x faster than SeqAn3’s FM-index implementation for nucleotide search, and ~2-6x faster for amino acid search in a single-threaded context. This performance is due to (i) a new approach to storing FM-index data in a strided bit-vector format that enables extremely efficient computation of the FM-index occurrence function via AVX2 bitwise instructions, and (ii) inclusion of a cache-efficient lookup table for partial k-mer searches. AWFM-index also trivially parallelizes to multiple threads with good scaling, and enables efficient on-disk storage of the memory-intensive suffix array. The open-source library is available for download at https://github.com/TravisWheelerLab/AvxWindowFmIndex.

## Background

String pattern matching is the problem of counting or locating occurrences of a query text pattern P within a large database text T. While not limited to the analysis of biological sequences, string pattern matching is integral to many tasks in bioinformatics, including mapping sequence reads to a reference genome [1, 2], taxonomic classification [3, 4], sequencing error correction [5], and seeding for sequence alignments [6, 7, 8].

The need for high-throughput pattern matching in bioinformatics has motivated myriad approaches including hashing, lookup tables, suffix arrays [9], and compressed suffix array data structures such as the FM-index [10]. Use of the FM-index across bioinformatic applications is due to its fast performance and low memory footprint. Unfortunately, its adoption is likely limited by the lack of an optimized and lightweight FM-index library; the only robust, currently maintained FM-index implementation we are aware of is found in the SeqAn3 library [11]. Here, we present a lightweight, open-source library called AvxWindowedFMindex (hereafter shortened to AWFM-index), which enables optimized string pattern matching over nucleotide or amino acid sequence datasets with significantly faster performance than SeqAn3’s library.

AWFM-index achieves significant performance gains through multiple algorithmic and data structure changes over a traditional FM-index implementation. Rather than storing the database text T in ascii symbols or as a range of integral values representing the symbols in T, AWFM-index stores bit-compressed symbols strided over 256-bit (AVX2) vectors that can be efficiently reduced with a low number of bitwise SIMD instructions. A table of k-mer seed ranges makes it possible to skip an early portion of the search computation for every query. Collections of multiple k-mers are queried in a thread-parallel manner, with good parallel scaling performance. AWFM-index is an open-source library written in C, with a simple API to facilitate easy integration into bioinformatics tools.

### Data Structure Background

#### Suffix Array

The suffix array [9] is a classic data structure that supports efficient determination of the count and locations of all occurrences of a query pattern P within a database sequence T. Given a text T that ends with a special sentinel symbol ‘$’ (defined as a symbol in the text’s alphabet Σ that otherwise does not occur in T, and is the smallest symbol in Σ), a suffix array SA is a permutation of integers [0..|T|-1], such that the suffix of T beginning at position SA[i] is lexicographically smaller than the suffix denoted by SA[j] if and only if i < j.

Because a suffix array lexicographically orders the suffixes of T, all indices of a given substring of T can be found in a contiguous range of elements in the suffix array. This fact is the key to the suffix array’s fast search, as it enables counting in O(|P| log|T|) time through binary search across the suffix array, and locating in O(|P| log|T| + k) time for k instances of the pattern. Without any data compression techniques, suffix arrays generally require 4 bytes of per symbol for sequences < 4GB long, or 8 bytes per symbol for sequences ≥ 4GB.

Numerous efficient algorithms have been devised to quickly construct a suffix array from text T. The optimal asymptotic performance for suffix array construction is O(|T|) [12], but the O(|T| log |T|) complexity divsufsort [13] is commonly used because of its excellent speed as an in-memory suffix sorter for genome-scale inputs; AWFM-index utilizes libdivsufsort [14] for suffix array construction.

#### Burrows-Wheeler Transform

The Burrows-Wheeler Transform (BWT) is a reversible text transform that was originally proposed for lossless data compression [15]. Given a text T and a associated suffix array SA, a BWT is defined as the transformation:

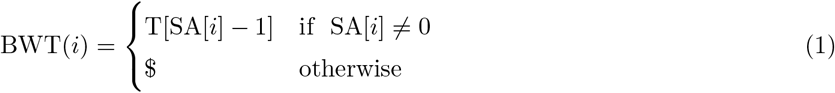

In other words, each element in the BWT holds the symbol directly preceding the suffix denoted at that element’s index in the suffix array. This is effectively the last column in a table of sorted rotations of T (see Fig 1), and is easily computed from a suffix array on T.

**Fig 1.**
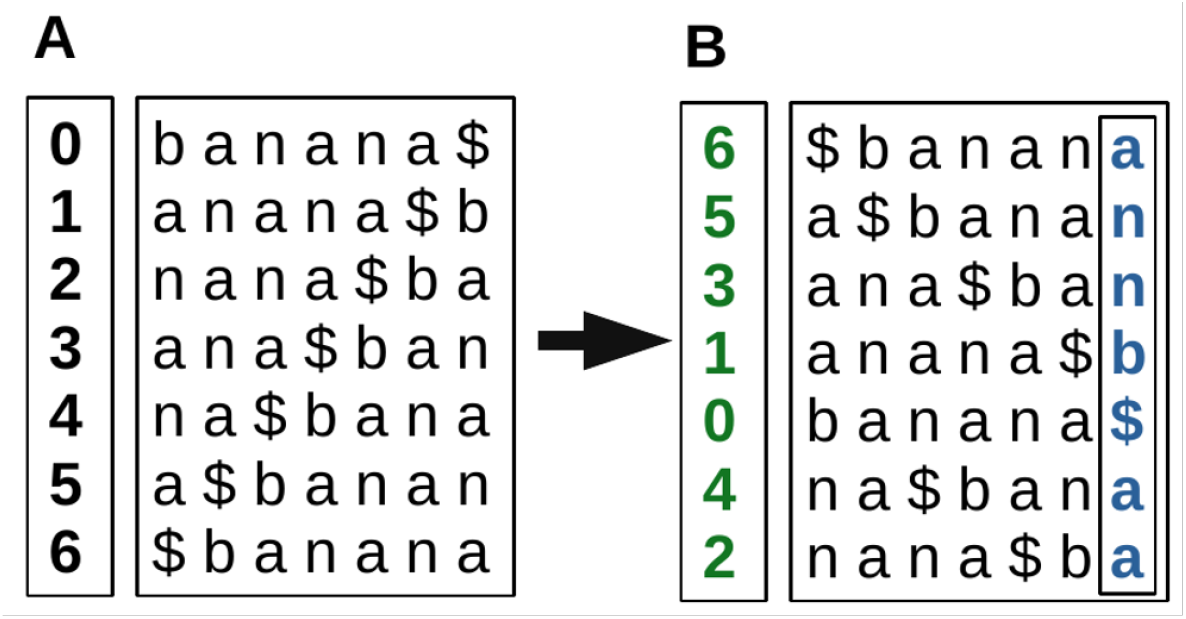
Example of generating a Burrows-Wheeler Transform for a given text. (A) All rotations of the input text ‘banana’, with appended sentinel ‘$’ symbol. The position of each rotation is given in the left column. (B) After sorting the rotations, the left column retains the original position of each rotation, and is thus the suffix array of the text. The final column of this sorted rotation matrix is the BWT. Note: the actual rotation matrix need not be stored, or even computed; it is represented here as a visual aid.

In order to reduce the memory footprint of a BWT, it is often losslessly compressed in some way. Strided bit vectors are commonly used as compressed BWT implementations, especially as a wavelet tree [16]. Wavelet trees are an attractive implementation, as they allow for lossless data compression approaching the empirical entropy of the text. For our implementation, we opted instead to use a new (and simpler) strided bit vector format that, along with precise symbol representations, enables efficient vector-parallel computation of the occurrence function.

#### FM-index

While the BWT can be viewed as a data-product of a suffix array, it can also be used as an alternative method for identifying pattern matches when used in conjunction with a suffix array. Using both data structures, Ferragina and Manzini introduced the FM-index [10]. An FM-index constructed from a given text T of alphabet Σ is comprised of the following: a suffix array SA, a Burrows-Wheeler Transform B, a milestone table described below, and a counts array C where C[s] is the count of all symbols in T that are lexicographically less than or equal to symbol s. Using these data structures, an FM-index can perform two key query functions called Count() and Locate(). The Count() function returns the number of occurrences of the query pattern P in *O*(|*P* |) time. The Locate() function returns the position in T of all k instances of P, in expected time *O*(|*P* | + *k*).

#### Exact pattern matching with FM-index

Search for a pattern P in text T is performed one character at a time, beginning with the final character of the pattern and moving backwards. To begin, the search process establishes a start-pointer and end-pointer [SP..EP] that correspond to the range in SA pointing to all occurrences of the final letter of P in T (Alg 1). In each successive step, the preceding character in P is prepended to the searched string P’, and the range [SP..EP] is updated (via Alg 2) to correspond to all positions in the text T that match the growing suffix, P’. This continues until each symbol in P has been processed. SP and EP are updated each time a new prefix symbol is added to the query using a function called occ() (short for occurrence). The occ(s, p) function takes as parameters a symbol *s* ∈ Σ − {$} and a position p where 0 ≤ *p* < |*T* |, and returns the number of occurrences of s in B before position p. To avoid unnecessary counting over large ranges of B, a milestone table is used to store the count S[s, p’] of symbol s preceding regularly sampled positions p’. When computing occ(s, p), the closest milestone position p’ < p is identified, and the value S[s, p’] is added to the count of symbol s between p’ and p. If the interval between milestones is r, the milestones table will require (Σ · ⌊|*B*|/*r*⌋ · 8) bytes, assuming 64-bit integers are used to store the symbol counts. After the conclusion of Alg 3, the range [SP..EP] of the query is the range of positions in the suffix array that represent suffixes that begin with pattern P (i.e., the locations of P). If at any point of the process SP > EP, P is not a substring of T, and the backward search is halted.

##### Algorithm 1

Constructing the initial [SP..EP] range for pattern P.

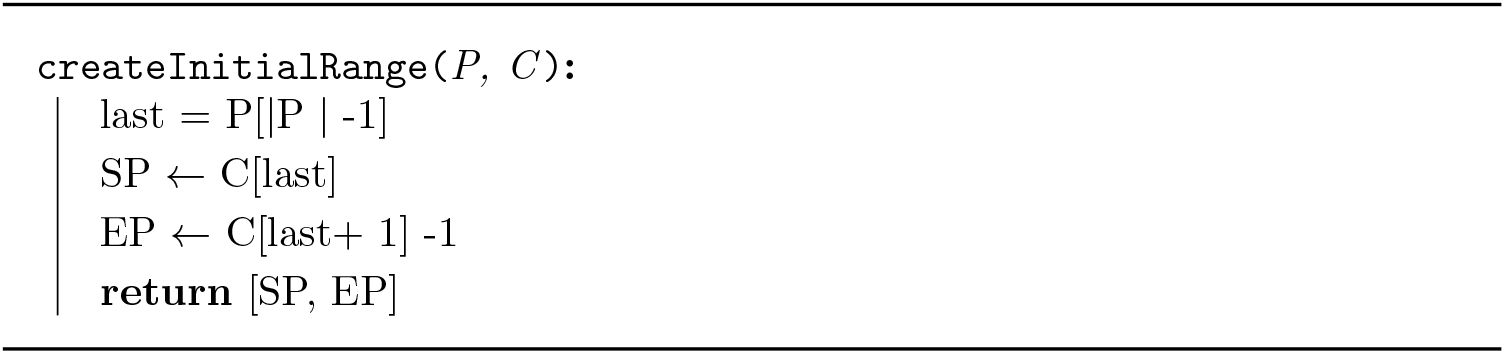

##### Algorithm 2

FM-Index backward search extending a BWT range [SP, EP] with prefix symbol s.

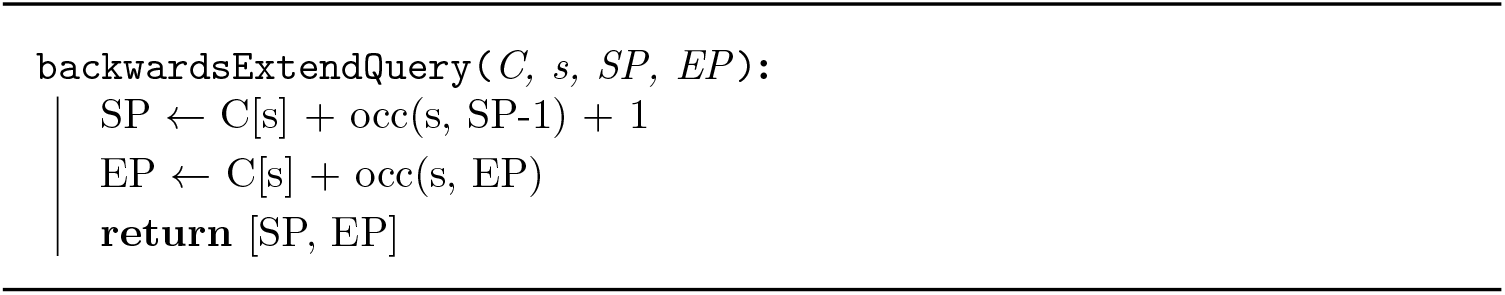

##### Algorithm 3

FM-index backwards search.

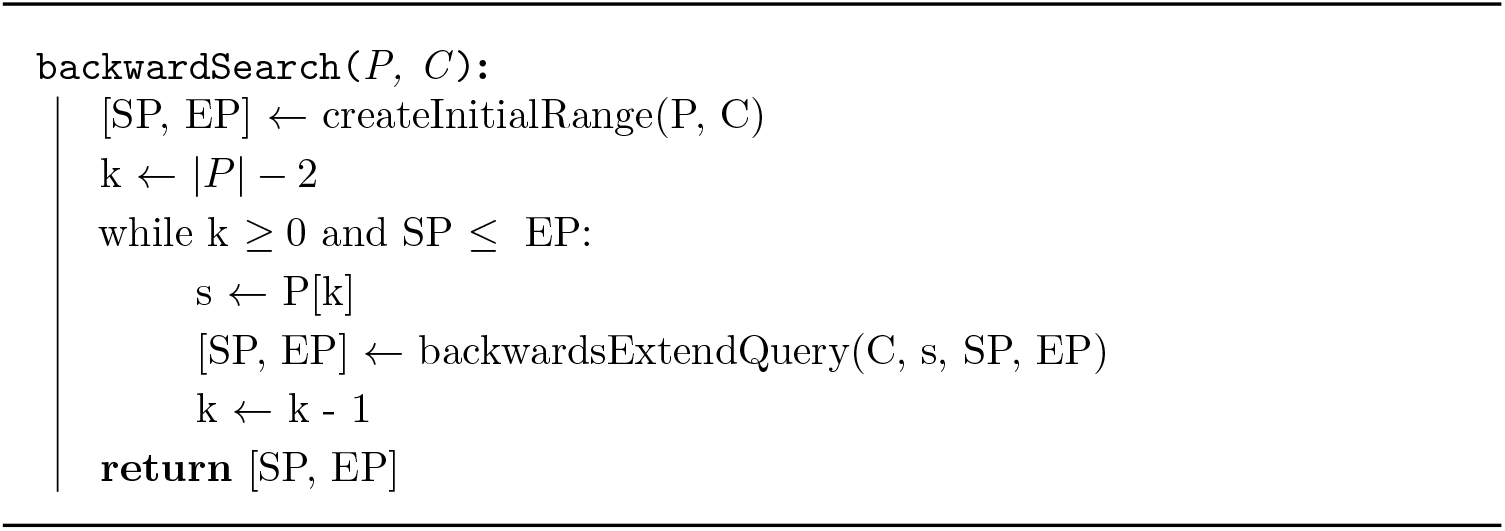

#### Reducing space requirements by sampling the suffix array

While locating query sequences using an FM-index has better complexity scaling than using only a suffix array for the same task (O(|P|) for FM-index as opposed to O(|P| log |T|) for suffix arrays), a naive FM-index requires more memory than suffix array alone, since it includes the BWT and milestone counts. By down-sampling the suffix array [17], the memory footprint of the suffix array inside an FM-index can be dramatically reduced at the cost of a modest performance hit. A common SA sampling strategy is called subscript sampling [18], in which a sampling ratio r is chosen, and the sampled SA’ is generated from all SA values at positions p where p ≡ 0 (mod r).

At the conclusion of the backwardSearch algorithm, each position in the [SP..EP] range corresponds to a position in the full SA, which itself indicates the location of an instance of P in T. Under SA down-sampling, only 1/r of the positions in [SP..EP] are present in SA’ (i.e., only positions p ≡ 0 (mod r) for p in [SP..EP]). For positions that are not sampled, the backtracePosition() function steps backwards through positions in the original text until a position sampled in SA’ is reached, then returns the correct position by adding the number of steps that were taken to this SA’ value.

A position p in B references some position B[p] in T. The backtrace step seeks to walk back in T until finding a position sampled by SA’; by construction, this is the character found at B[p]. Thus, to take one step back in T from a current position p, the symbol at p in B is found and the symbol is used in the occurrence function to find the BWT position of the previous symbol in the original text T (Alg 4).

##### Algorithm 4

Backtracing a position p to the nearest previous position sampled in SA’; this supports identification of the position in T corresponding to position p in SA.

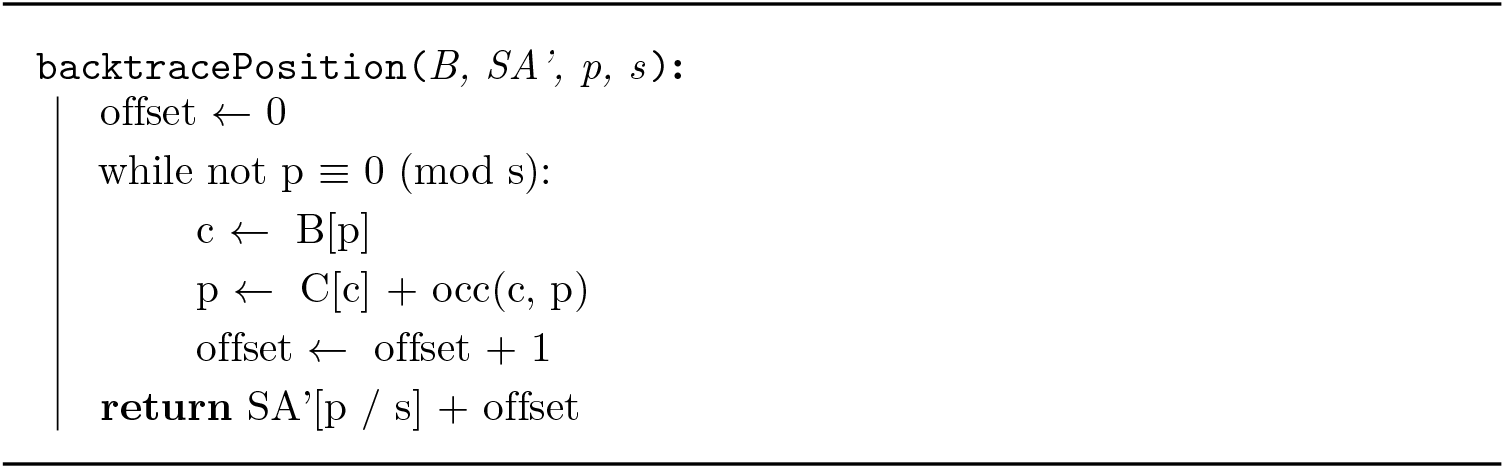

## Implementation

This manuscript describes an optimized FM-index library that is lightweight, easy-to-incorporate, and provides clients with FM-index functionality at both a high-level (count or locate all instances of a query string) and low-level (step-wise control of the FM-index backward-search process). Here, we present the various strategies that contribute to the library’s fast text indexing performance. The key innovation is the development of a representation of BWT sequence data with a strided bit-compressed vector format; this is interleaved with milestone data in a manner similar to [19]. This format supports efficient computation of the most expensive aspect of FM-index calculations: the occurrence function.

We begin by describing a specialized bit representation for symbols in both nucleotide and amino acid alphabets, along with an efficient method testing symbol equality with such a representation. We then show how this symbol representation can be used to compactly store an FM index in memory blocks representing 256 symbols at a time, and that these blocks can be efficiently processed using AVX2 vector instructions. This is followed by description of other aspects of the implementation, including a partial k-mer query lookup table that allows AWFM-index to skip the first few [SP..EP] update steps for each query.

### Bitwise Symbol Matching

Consider an alphabet Σ, with each symbol in the alphabet encoded using n bits. In order to count the occurrences of a query symbol s in a range of symbols in the BWT, each symbol in the range must be checked for equality to s. While nearly all CPU architectures contain instructions to directly compare two numbers, we explore solutions that exploit bitwise operations for comparing symbols. Given two symbols *s*_1_, *s*_2_ ∈ Σ, one simple method for comparing *s*_1_ against *s*_2_ is to use a straightforward combination of bitwise operations: (i) for all set bits in *s*_1_, the corresponding bits in *s*_2_ are ANDed together; (ii) for all clear bits in *s*_1_, the corresponding bits in *s*_2_ are ORed together, and then bitwise NOTed. The boolean values that result from these actions are then ANDed together. The resulting value is true iff *s*_1_ = *s*_2_, and can be computed in n bitwise operations, or n-1 operations for the case where all bits in *s*_2_ are set. By performing bitwise operations in this way, symbol equality can be checked even in situations where a direct symbol comparison operation is not possible (as is the case when performing vectorized computations, as described shortly). For the purposes of this implementation we consider the following bitwise operations on a and b: AND(a,b) = a&b, OR(a,b) = a|b, and ANDNOT(a,b) = (!a)&b to each be single bitwise operations, as they are each a single CPU instruction within our target instruction set.

In AWFM-index, two alphabets are supported, one for nucleotide data, one for amino acid data. Each alphabet contains symbols for each of the possible residues (4 for nucleotides, 20 for amino acids), a sentinel symbol, and an ambiguity symbol, denoted here as X, defined to be lexicographically greater than all other symbols in Σ. The resulting alphabets are length 6 and 22 respectively, and symbols in each alphabet are represented with ⌈log_2_(6)⌉=3 and ⌈log_2_(22)⌉=5 bits. Note that each of these alphabets have fewer symbols than the number of possible values for each of their corresponding bit lengths. A naive approach to assigning encodings to the |Σ| symbols in each alphabet would be to assign them to the integers [0 .. |Σ| − 1]. Instead, AWFM-index assigns alphabet symbol encodings using a strategy that aims to reduce the number of bitwise operations needed to compare symbols for equality. These encodings are presented in Table 1, and explained in the next two sections.

**Table 1.** **Bit encodings** for all nucleotide and amino acid symbols, and the number of bitwise operations required to check for equality when used as a query symbol.

| Nucleotide<br>IUPAC Code | Bit Encoding | Symbol Group | # Bitwise Ops<br>for Comparison |
| --- | --- | --- | --- |
| A | 110 | 1 | 1 |
| G | 101 |  |  |
| C | 011 |  |  |
| T | 001 | 2 | 2 |
| X | 010 |  |  |
| \$ | 100 | | |
| Amino Acid<br>IUPAC Code | Bit Encoding | Symbol Group | # Bitwise Ops<br>for Comparison |
| A | 01100 | 1 | 2 |
| D | 00011 |  |  |
| E | 00110 |  |  |
| G | 11010 |  |  |
| I | 11001 |  |  |
| K | 11001 |  |  |
| L | 11100 |  |  |
| P | 01001 |  |  |
| R | 10011 |  |  |
| S | 01010 |  |  |
| T | 00101 |  |  |
| V | 10110 |  |  |
| C | 10111 | 2 | 3 |
| F | 11110 |  |  |
| H | 11011 |  |  |
| M | 11101 |  |  |
| N | 01000 |  |  |
| Q | 00100 |  |  |
| W | 00001 |  |  |
| Y | 00010 |  |  |
| X | 11111 | 3 | 3 |
| \$ | 00000 | | |

#### Nucleotide alphabet symbol encodings

Nucleotide symbols are represented by two groups of unique 3-bit encodings. Group-1 encodings have 2 of the 3 bits set, while group-2 encodings have only a single set bit. With a group-1 nucleotide query symbol, equality to another symbol is determined by ANDing the 2 bits corresponding to the set bits of the query. This produces a true boolean result if the symbol matches, and precludes a true result for any other symbol: (i) any other group 1 encoding would have a different pair of set bits, so that one of the compared bits would not be set, yielding a false result from the AND operation; (ii) any group-2 symbol contains only one set bit, so again the AND operation would return false, and (iii) since no encodings have more than 2 bits set, we can be sure that no other symbol could match to our query. Group-2 encodings can be checked for equality in 2 bitwise operations by taking the ANDNOT of the set bit and one of the clear bits, then ANDNOTing the result with the last clear bit. This strictly forces each of the 3 bits to match the query symbol.

#### Amino acid alphabet symbol encodings

Amino acid symbol encodings are split between 3 groups of unique 5-bit encodings. Group-1 encodings are represented by all 12 possible encodings in which exactly 2 of the lower 4 bits (bits [0..3]) differ from the most significant bit (bit 4). Group-2 encodings are represented by all 8 possible encodings in which exactly 1 of the bits in [0..3] differs from bit 4. Group-3 encodings are represented by all 5 bits being either set or cleared, and denote the ambiguity symbol and the sentinel respectively. For a group-1 amino acid query symbol, equality can be tested in 2 bitwise operations, with the required operations depending on the state of bit 4 in the query symbol. If a query symbol is in group-1 and its bit 4 is set, one of the two bits corresponding to the query’s clear bits is ANDNOTed with bit 4, then the other clear bit is ANDNOTed with the result. If bit 4 is clear for a group-1 symbol, the two bits corresponding to the query set bits are ANDed together, and the result is ANDNOTed with bit 4. Both of these options return true if and only if bit 4 matches the query, and the 2 bits that are supposed to differ from bit 4 in fact do so. Further, if the result is true, it shows that the symbol cannot be a group-2 encoding, since more than 1 bit differs from bit 4. In the same way, it shows that the symbol cannot encode for group-3. If the result is true, therefore, it cannot encode for any symbols other than our query. For a group-2 amino acid query symbol, equality can be tested in 3 bitwise operations. If bit 4 is set, the bit corresponding to the query’s single clear bit is ANDNOTed with one of the 3 set bits in [0..3]. The other 2 set bits in [0..3] are ANDed together, and the result is ANDed with the result of the first operation. If bit 4 is clear, one of the 3 clear bits in [0..3] is ANDNOTed with the single set bit. The remaining 2 clear bits are ORed together, and the result is ANDNOTed with the result of the first operation. Note that we did not check bit 4 in any way; if the result is true we can infer the state of bit 4 because no encodings exist with 3 bits that differ from bit 4. Therefore the state of bit 4 must be the opposite of the bit with the unique state. We also know that the symbol cannot encode for a group-1 or group-3 symbol, because we have shown that exactly 1 bit differs from bit 4. Therefore, the result is true if and only if the symbol matches the query.

Group-3 comparisons are straightforward. The ambiguity symbol is encoded with 5 set bits, and can be compared by ANDing together bits 0 and 1, ANDing bits 2 and 3, and ANDing the two results. Again, bit 4 does not need to be checked because no symbols are represented by all bits in [0..3] differing from bit 4. Comparison against the sentinel symbol is similar, however this is never necessary in practice since query strings cannot contain the sentinel.

As group-1 encodings require 1 less instruction to reduce, this group is used to encode for the 12 most frequent amino acids in the UniProtKB/Swiss-Prot database [20] and group-2 encodings represent the 8 least frequent amino acids. The reason for this choice is that the more-common amino acids will likely be queried more often, and therefore should be represented by encodings that require the fewest instructions.

### Strided Bit Vector Data Format

The previous section described a bitwise method for comparing a single symbol encoding against a query symbol. AWFM-index employs this bitwise comparison strategy in the context of Single Instruction, Multiple Data (SIMD) parallelization. Specifically, AWFM-index uses AVX2 instructions to perform the bitwise operations on vectors of 256 symbols in parallel, effectively comparing up to 256 symbols from the BWT to a single query symbol in the same number of bitwise instructions as comparing a single symbol. The AVX2 instruction set is an extension of the x86 instruction set that performs operations on vectors of 256 bits with a single instruction. AWFM-index uses 3 AVX2 intrinsic instructions (mm256 and si256, mm256 or si256, and mm256 andnot si256) to implement the bitwise operations for comparing symbol encodings, as described earlier. Through precise, interleaved layout of the BWT and milestone data, AWFM-index finds all matches to a query symbol in ≤4 instructions, and computes the occurrence function with a few extra steps.

In the AWFM-index, the BWT sequence is broken up into windows of 256 symbols; each window represents a range [i..j] (with j=i+255), and is comprised of 3 sections: (i) multiple contiguous 256-bit AVX2 vectors that store the 256 symbols in a strided bit-vector format (see Fig 2, and text below), (ii) an array of 8-byte milestone occurrence counts containing the count of symbol s in B[0..i-1], for each symbol in the alphabet except the sentinel, and (iii) a padding section to ensure that all strided bit vectors align to 32-byte boundaries necessary for AVX2 SIMD instructions. By interleaving the milestone counts with the BWT data, both the milestone count and the symbol bit vectors can be brought into cache in the same memory request. The milestone section of the window contains five (5) 8-byte values for nucleotide windows, or twenty-one (21) 8-byte values for amino acid windows. Since the milestone sections are aligned to a 32 byte (256 bit) memory boundary, a 24 byte padding (for both nucleotide and amino acid alphabets) ensures that the strided bit-vector section also aligns to a 32 byte boundary.

**Fig 2.**
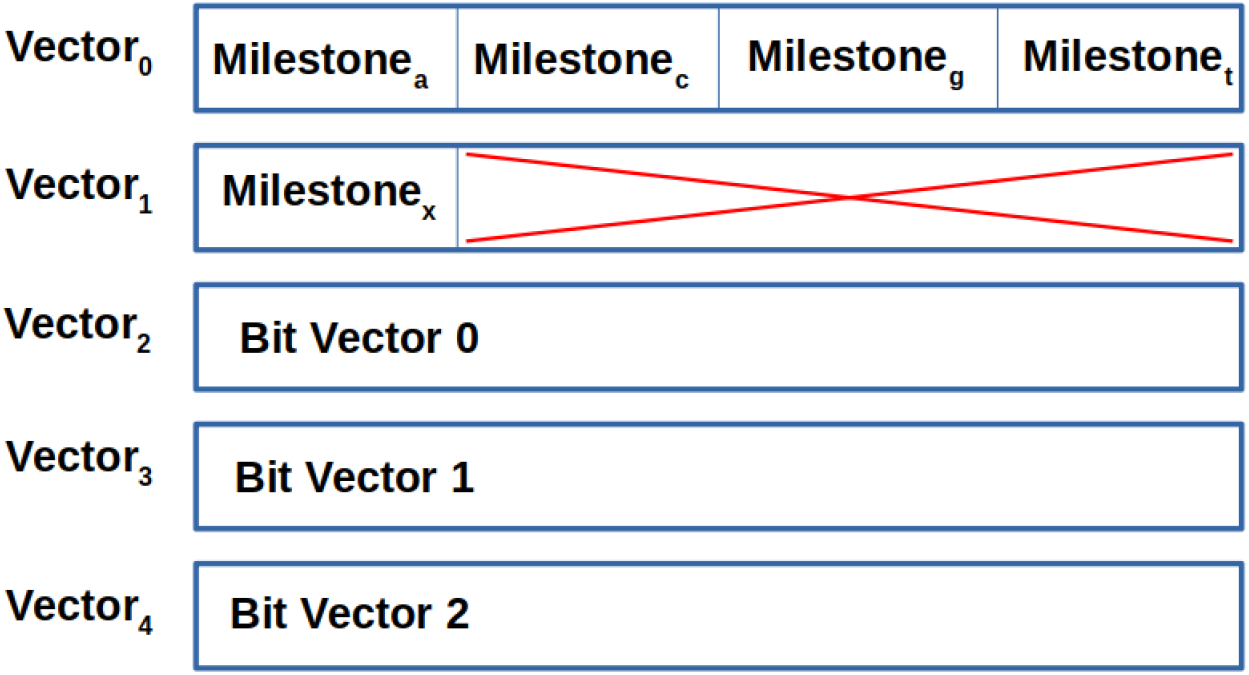
The 5 AVX2 vectors that comprise a nucleotide BWT window. The milestone counts for A, C, G, and T are stored in vector 0. Vector 1 contains the milestone for the ambiguity symbol ‘X’ and a 24 byte padding section to align the bit vectors to the 32 byte alignment necessary for AVX2 instructions. Vectors 2-4 contain the bits representing the symbols in the window.

As a BWT window denotes a contiguous range of 256 symbols from the BWT, a straightforward approach to storing symbols would be to represent each symbol with one byte as raw ASCII character values. For small alphabets, modern FM-index implementations prefer to use some form of bit-compressed representation, such as representing nucleotide symbols with 2 bits [1] (though this approach does not support ambiguity symbols, and special handling is required for the sentinel symbol). AWFM-index adjusts this bit-compression strategy to better leverage SIMD parallelization in computing the occurrence function. The symbols in the BWT are strided over the window’s bit-vectors, with one vector for bit-0 of all 256 symbols, another vector for bit-1, and so on (generally: bit n of bit-vector m of a given window represents the m-th bit of the n-th symbol in the window). Including the milestone values and the padding, nucleotide data takes up 5 AVX2 vectors for each 256 symbol window, and thus requires 5 bits per symbol in the original text. Similarly, amino acid windows take up 11 AVX2 vectors, and so require 11 bits per symbol.

### SIMD Occurrence Calculation

To compute the occurrence function occ(s, p) for symbol s and position p, the milestone occurrence count is taken from the appropriate section in the BWT window. Then, positions in the BWT window matching symbol s are identified and captured into a 256-bit vector such that bit n is set if and only if the n-th position in the window represents symbol s, here called an occurrence vector. This occurrence vector is generated by using AVX2 SIMD instructions to implement a bitwise comparison across all 256 symbols in the window, with bitwise instructions described in the previous section (see Fig 3). Once the occurrence vector has been generated, a bitmask is applied to clear all bits after position p; this ensures that no positions after p are counted in the final occurrence count. The set bits in each of the 4 quad-words in the occurrence vector are then counted with popcnt64() intrinsic instructions, and the results are summed with the corresponding milestone count to compute the final occurrence count. Multiple strategies of generating a population count of the occurrence vector were tested, and summing the results of the 4 popcnt64() instructions was found to outperform other SIMD vector popcounting techniques (e.g. [21]).

**Fig 3.**
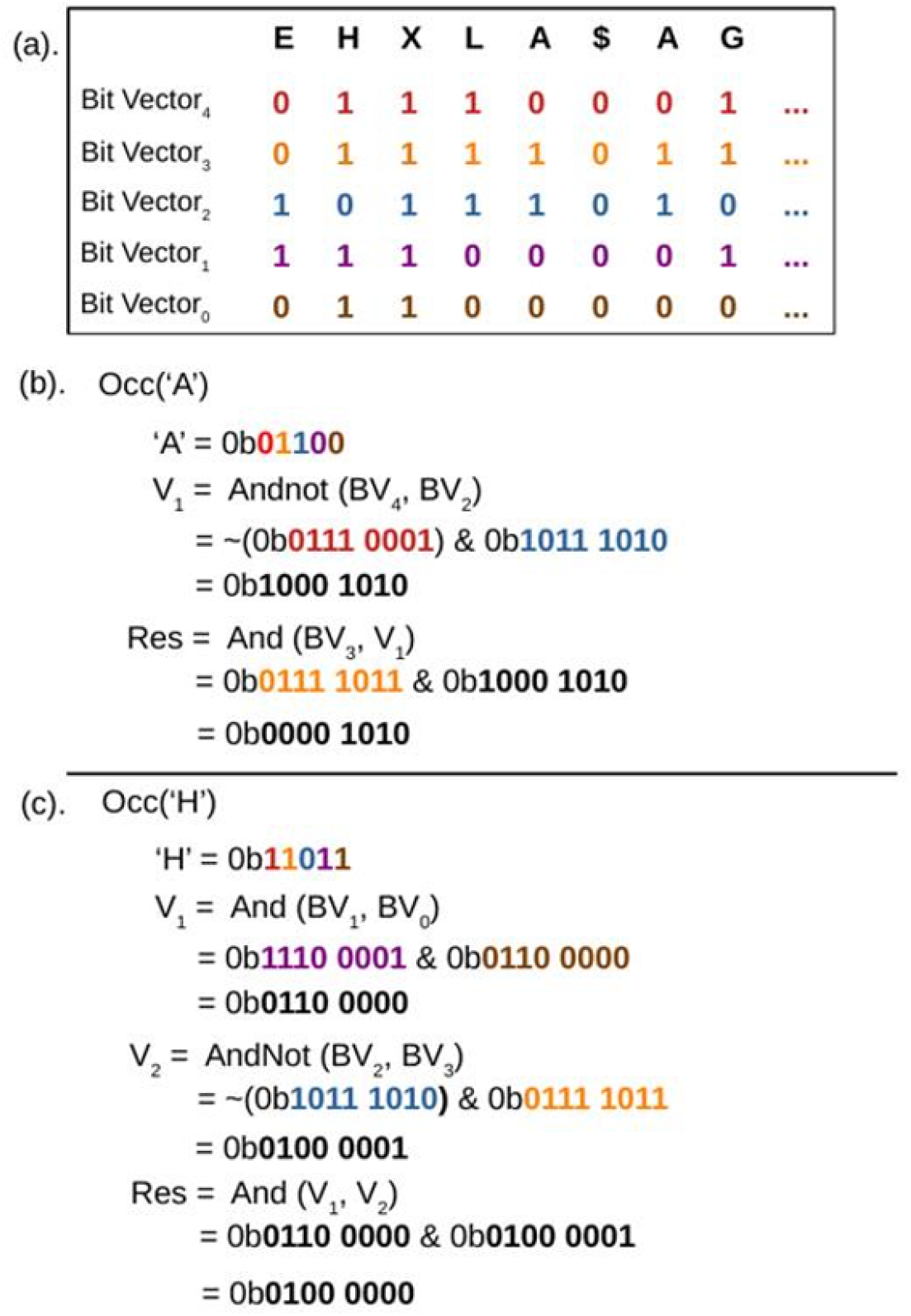
Examples of creating an occurrence bit vector from the strided BWT bit vectors. (a) An example of the bit vectors in a BWT window. Each bit vector is 256 bits wide representing 256 symbols, but only the first 8 positions are shown for brevity. By performing bitwise operations on these bit vectors an occurrence vector can be generated where a set bit indicates the presence of the queried symbol at the bit’s position in the window. (b) Creating an occurrence bit vector for Group 1 amino acid ‘A’ in 2 SIMD operations. (c) Creating an occurrence bit vector for Group 2 amino acid ‘H’ in 3 SIMD operations.

### Manual Prefetching

While the computation to update the BWT range is minimal, the unpredictable nature of each subsequent position p given to the occurrence function creates a performance bottleneck in reading data from memory. Given a sufficiently large BWT, every occurrence call will result in a memory read request for cache lines that will almost certainly not be in cache, except for pathological case queries like ‘AAAAAAA’, or if SP and EP land in the same BWT window. To ease the performance hit caused by this random access, AWFM-index employs manual data prefetching using the _mm_prefetch() SSE instruction as soon as the location of the memory address for the next occurrence function is known. The updates to SP and EP are also staggered such that after SP is updated, a prefetch request is generated for the following SP, and the update to EP begins while the new SP memory prefetch request is being serviced.

### Accelerated Search with a K-mer Lookup Table

Traditionally, searching an FM-index involves updating the [SP,EP] pair for every symbol in the given query. AWFM-index uses a pre-computed lookup table with a modest memory footprint to skip a sizable portion these [SP,EP] updates. When the AWFM-index is built, a parameter k is selected, and a table is allocated to store the [SP,EP] pair for every length-k string over alphabet Σ (except the ambiguity and sentinel symbols, which are excluded because neither are found in query strings). This table of k-mer ranges consumes 16 · (|Σ| − 2)^*k*^ bytes in memory, and memoizing prefix ranges enables *O*((|Σ| − 2)^*k*^) construction time. If a query pattern P is at least length k, the [SP,EP] range for the length-k suffix of P is found in the k-mer lookup table, effectively skipping the first k updates to the [SP,EP] range. Then, the query proceeds as normal, querying for symbols until the entire query has been completed, or the range is invalid. If the query string length is less than k, the range is resolved without using the k-mer table. The recommended values of k are 12 for nucleotide indices (268MB table size) and 5 for amino indices (51MB table size) as they strike a balance between memory footprint and performance benefit. Other values may be selected by the library client depending on expected factors such as expected query lengths and available system memory.

### API and Thread-Parallel Search

The core API for AWFM-index includes the locate() and count() functions, which each accept as arguments (i) the AWFM-index data structure, (ii) a collection of query sequences, and (iii) a number of threads used to parallelize search. Parallelization is achieved using simple OpenMP 1.0 pragmas, as each query in the collection is data-independent with respect to the other queries in the collection. Given a collection Q of query sequences, the collection is implicitly divided into batches of 4 queries, resulting in a total of ⌈|Q| / 4⌉ batches. The user-specified number of threads are then used to parallelize the search across the collection of batches. When a thread begins to compute the results for a batch of queries, it begins by finding [SP,EP] range in the k-mer lookup table that represents the final k symbol suffix for each of the 4 queries. Then, each query in the batch is extended until either the SA range of the query has been fully resolved, or has failed due to SP > EP. If the parallel locate() function is used, the location of each instance of each query string is found via the SA backtracking method described earlier. We chose 4 for the batch size so that each thread can work on a group of contiguous queries and to hide the cost of thread management, but not such a large batch that cache eviction becomes a performance concern. We tried multiple values for the batch size, but since most small batch sizes performed similarly, 4 was an essentially arbitrary choice.

The AWFM-index API also includes non-parallelized functions for initializing a SA range, extending queries with additional prefix symbols, backtracing to the most recently sampled SA position, and looking up the original position using the suffix array. These functions allow a client to implement custom FM-index applications based on the internal components of AWFM-index, for example for inexact pattern matching.

### Suffix Array Sampling, In-Memory or On-Disk

The suffix array component of the FM-index is often down-sampled to reduce memory requirements. Suffix arrays that are sparsely sampled have a smaller memory footprint, but require more backtrace steps to deduce the actual sequence position during the locate() function. The AWFM-index library currently supports suffix array sampling ratios r that are 1 ≤ r < 256 and utilizes a subscript sampling strategy such that every r^*th*^ entry is sampled. AWFM-index provides the option to either load the suffix array into memory with the rest of the index (default) or leave the suffix array on disk and read directly from disk the values necessary to resolve the final sequence positions after all SA range elements have been backtraced to a sampled SA position. While disk access is significantly slower than memory access, on-disk suffix array storage allows for denser sampling, even on systems with limited memory. For example, in the context of the locate() function, using an AWFM-index with a suffix array sampling ratio of 1 where the suffix array is left on disk results in a single random disk read for each found substring, which may be preferable to the large number of sequential cache misses necessary to backtrace to the nearest sampled suffix array position for each found substring in a heavily downsampled suffix array. This functionality aims to make high performance indexing accessible to a wider range of users on personal computers or laptops with limited memory.

### Suffix Array Minimum Bit-Width Compression

To reduce memory requirements, AWFM-index stores SA values as variable bit-width integers similar to the int vector class of SDSL[22], rather than as simple 64-bit integers. Given a suffix array S of length n, all values within S are non-negative and less than n. Therefore, each value can be represented with ⌈log_2_(*n* − 1)⌋ bits. Each sample in the suffix array is compressed to this many bits, and repacked into a byte array. An individual value can then be extracted in constant time back into a 64-bit integer. Storing the suffix array in this minimum bit-width integer array results in a reduction in suffix array space requirement of ⌈log_2_(|T| + 1)⌉·⌊(|T|+1)/r⌋ bits for compression ratio *r*.

## Results

We performed numerous tests to assess the performance of AWFM-index relative to the FM-index implementation inside SeqAn 3.0.3 [11], and to demonstrate the impact of AWFM-index parameterization. Unless stated otherwise, All tests were run on a system with a 32-core Intel Xeon E5-2630 v3 @ 2.40GHz, and 64 Gb RAM.

### Speed Comparison Between AWFM-index and SeqAn3

A 1 billion base pair nucleotide sequence and a 200 million amino acid sequence were generated with the easel sequence analysis library [23]. FM-index files were generated from these sequences for both SeqAn3 and AWFM-index using the native construction method for each tool. The index build times were similar for the two libraries. A partial lookup table was pre-computed for all length-12 (nucleotide) and length-5 (amino acid) k-mers. A collection of 1 million queries of varying lengths were sampled from the original text, and run times for locate() and count() functions were captured in Tables 2 and 3. Count() calls were typically 2-6x faster with AWFM-index, while Locate() calls were typically 2-4x faster.

**Table 2.** Run time for nucleotide locate() and count() function. Target consists of 1 billion nucleotide-long simulated sequence with a suffix array sampling ratio of 4. Query consists of 1 million nucleotide query sequences sampled from the target at each of several query lengths.

| Nucleotide search |  | Count()<br>Time (seconds) |  |  | Locate()<br>Time (seconds) |  |  |
| --- | --- | --- | --- | --- | --- | --- | --- |
| Query Length | Hits/Query | SeqAn3 | AWFM | Speed-up | SeqAn3 | AWFM | Speed-up |
| 20 | 1.00 | 4.51 | 1.15 | 3.12 | 6.14 | 2.37 | 2.60 |
| 18 | 1.01 | 3.96 | .97 | 4.07 | 6.93 | 1.72 | 4.04 |
| 16 | 1.23 | 3.50 | 1.06 | 3.29 | 5.21 | 2.57 | 2.02 |
| 14 | 4.73 | 2.90 | .76 | 3.80 | 8.61 | 3.67 | 2.34 |
| 12 | 60.60 | 2.85 | .18 | 15.85 | 42.46 | 29.65 | 1.43 |
| 11 | 239.40 | 1.87 | .99 | 1.90 | 147.54 | 106.51 | 1.39 |

**Table 3.** Run time for amino acid locate() and count() functions. Target consists of 200 million character-long simulated amino acid sequence with a suffix array sampling ratio of 4. Query consists of 1 million amino acid query sequences sampled from the target at each of several query lengths.

| Amino acid search |  | Count()<br>Time (seconds) |  |  | Locate()<br>Time (seconds) |  |  |
| --- | --- | --- | --- | --- | --- | --- | --- |
| Query Length | Hits/Query | SeqAn3 | AWFM | Speed-up | SeqAn3 | AWFM | Speed-up |
| 10 | 1.00 | 4.58 | .81 | 5.66 | 6.40 | 1.63 | 3.92 |
| 9 | 1.00 | 3.91 | .83 | 4.71 | 5.97 | 1.38 | 4.33 |
| 8 | 1.02 | 3.39 | .60 | 5.69 | 5.29 | 1.46 | 3.61 |
| 7 | 1.47 | 2.89 | .43 | 6.77 | 5.41 | 1.61 | 3.36 |
| 6 | 9.00 | 2.38 | .38 | 6.27 | 15.35 | 5.52 | 2.78 |
| 5 | 137.70 | 1.81 | .08 | 21.95 | 167.56 | 73.42 | 2.28 |

### Effect of K-mer Lookup Table on Speed

To gauge the performance gains due to the partial k-mer lookup table, the previous nucleotide benchmark was used to compare AWFM-index locate() performance with a minimum size lookup table (k=1) versus the default recommended size partial k-mer look table (k=12). Fig 4 shows that count() performance is significantly boosted by avoiding the first 12 steps of Alg 3. Meanwhile the impact on locate() is modest, as large numbers of backtrace operations cause runtime to be dominated by occurrence calculations during backtrace.

**Fig 4.**
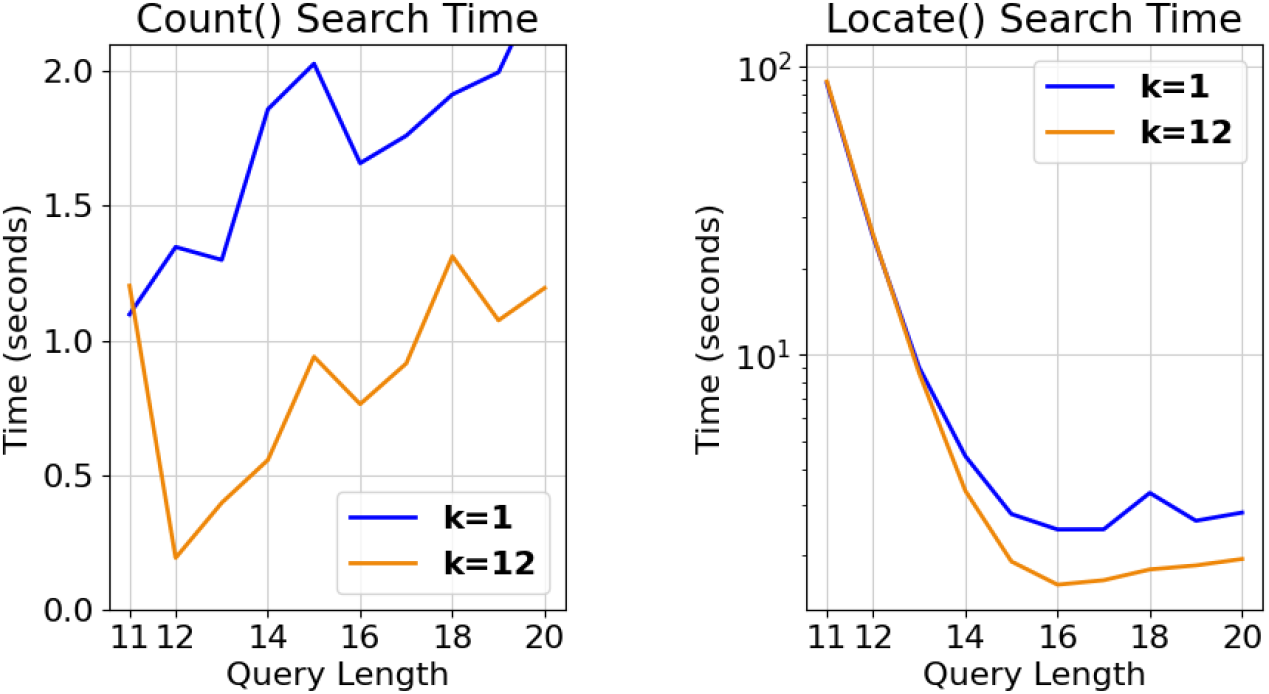
Timings of 1 million nucleotide queries using a partial k-mer table of length 1 (blue), and of length 12 (orange). A suffix array compression ratio of 4 was used for each index. The performance benefits of the partial k-mer table are most obvious in the count() function, whereas the performance benefit in the locate() function is most notable for longer queries that generate fewer hits (i.e. when the number of backtrace steps is relatively small).

### Memory Footprint and Suffix Array Sampling

To determine the performance characteristics of working with the suffix array on-disk, benchmarks were performed with SA kept either in-memory, on a hard-disk drive (HDD), or on a solid-state drive (SSD). These benchmarks were performed on a system with a Intel(R) Xeon(R) CPU E5-2620 v4 @ 2.10GHz processor. Not surprisingly, the performance loss by storing the suffix array on-disk varies depending on whether disk storage uses hard disk drives or solid state drives. When stored on solid state drives, fully-sampled on-disk suffix arrays outperform in-memory suffix arrays at suffix array compression ratios of approximately 4, while generating a smaller memory footprint (Table 4). At higher compression ratios, the difference in performance between in-memory and on-SSD indices becomes negligible, since the time spent backtracing largely exceeds suffix array lookups. When an on-disk SA is stored on a HDD, the fully-sampled SA performs about as well as an in-memory SA with a compression ratio of 16; similarly to the SSD tests, the difference between on-disk and in-memory shrinks as the suffix array compression ratio increases. Since no memory is used for an on-disk SA, this provides an efficient mechanism for decreasing memory load while retaining speed, particularly if SSD storage is available.

**Table 4.** Impact of suffix array compression. Suffix array memory requirements for various suffix array compression ratios (target length = 1 billion), and the time taken to locate() 1 million length-14 nucleotide queries for in-memory and on-disk suffix arrays. The average number of hits per query was 4.73.

| Suffix Array |  | Locate() time (seconds) |  |  |
| --- | --- | --- | --- | --- |
| Compression ratio | Suffix Array size | In-memory SA | SSD SA | HDD SA |
| 1 | 3750 MB | 0.67 | 2.18 | 8.71 |
| 2 | 1875 MB | 1.28 | 3.44 | 6.18 |
| 4 | 938 MB | 2.41 | 4.75 | 6.64 |
| 8 | 469 MB | 4.72 | 6.98 | 8.40 |
| 16 | 234 MB | 9.42 | 12.34 | 11.98 |
| 32 | 117 MB | 19.00 | 20.66 | 21.10 |
| 64 | 59 MB | 37.07 | 39.42 | 39.58 |
| 128 | 29 MB | 75.25 | 77.14 | 79.25 |

Tables 5 and 6 compare the performance and memory usage of AWFM (in-memory) and SeqAn3 indices over a range of SA compression ratios. The SeqAn3 BWT implementation uses wavelet trees to store the BWT, resulting in a reduced memory footprint compared to AWFM. At more densely-sampled suffix arrays, the memory footprint differences are negligible compared to the performance gains over SeqAn3. With sparsely sampled suffix arrays, the BWT makes up a large fraction of the stored data structure, so that AWFM’s speed gains are offset by an increased memory requirement. Though SeqAn’s memory usage is lower than AWFM’s in-memory variant for any fixed SA compression ratio, AWFM is generally faster for any given memory footprint. Leaving the SA on SSD disk will improve this speed-vs-memory trade-off in AWFM’s favor.

**Table 5.** Comparing AWFM and SeqAn3 Locate() performance for nucleotide search, across various suffix array compression ratios. The target sequence is a 1 billion-character long nucleotide sequence generated by the easel tool ‘esl-shuffle’. Query consists of 1 million nucleotide queries of length 14 taken from the target sequence. Default k-mer lookup table size of 12 was used. Time and memory were captured with /usr/bin/time.

| Compression Ratio | Nucleotide Locate() - 1Gb |  |  |  |  |
| --- | --- | --- | --- | --- | --- |
|  | Time (seconds) |  |  | Memory (Mb) |  |
|  | SeqAn3 | AWFM | Speed-up | SeqAn3 | AWFM |
| 1 | 4.30 | 1.07 | 4.03 | 4007 | 4535 |
| 2 | 5.20 | 4.17 | 1.24 | 2176 | 2704 |
| 4 | 8.62 | 4.60 | 1.87 | 1261 | 1789 |
| 8 | 13.78 | 6.26 | 2.20 | 803 | 1331 |
| 16 | 24.48 | 11.90 | 2.06 | 574 | 1102 |

**Table 6.** Comparing AWFM and SeqAn3 Locate() performance for nucleotide search, across various suffix array compression ratios. The target sequence is a 200 million-character long amino acid sequence generated by the easel tool ‘esl-shuffle’. Query consists of 1 million amino acid queries of length 6 taken from the target sequence. Default k-mer lookup table size of 5 was used. Time and memory were captured with /usr/bin/time.

| Compression Ratio | Amino Acid Locate() - 200Mb |  |  |  |  |
| --- | --- | --- | --- | --- | --- |
|  | Time (seconds) |  |  | Memory (Mb) |  |
|  | SeqAn3 | AWFM | Speed-up | SeqAn3 | AWFM |
| 1 | 3.30 | 0.83 | 3.96 | 822 | 1003 |
| 2 | 7.05 | 2.60 | 2.71 | 481 | 661 |
| 4 | 15.37 | 6.21 | 2.47 | 310 | 490 |
| 8 | 34.33 | 12.95 | 2.65 | 224 | 405 |
| 16 | 79.01 | 27.49 | 2.87 | 182 | 362 |

### Thread-Parallel Performance

We evaluated the speed gains achieved with multi-threading using the nucleotide benchmark described above (1 billion simulated nucleotides), with 1 million length-14 query strings. As seen in Fig 5, AWFM-index presents ∼35-50% strong scaling efficiency up to 20 threads, with diminished returns thereafter.

**Fig 5.**
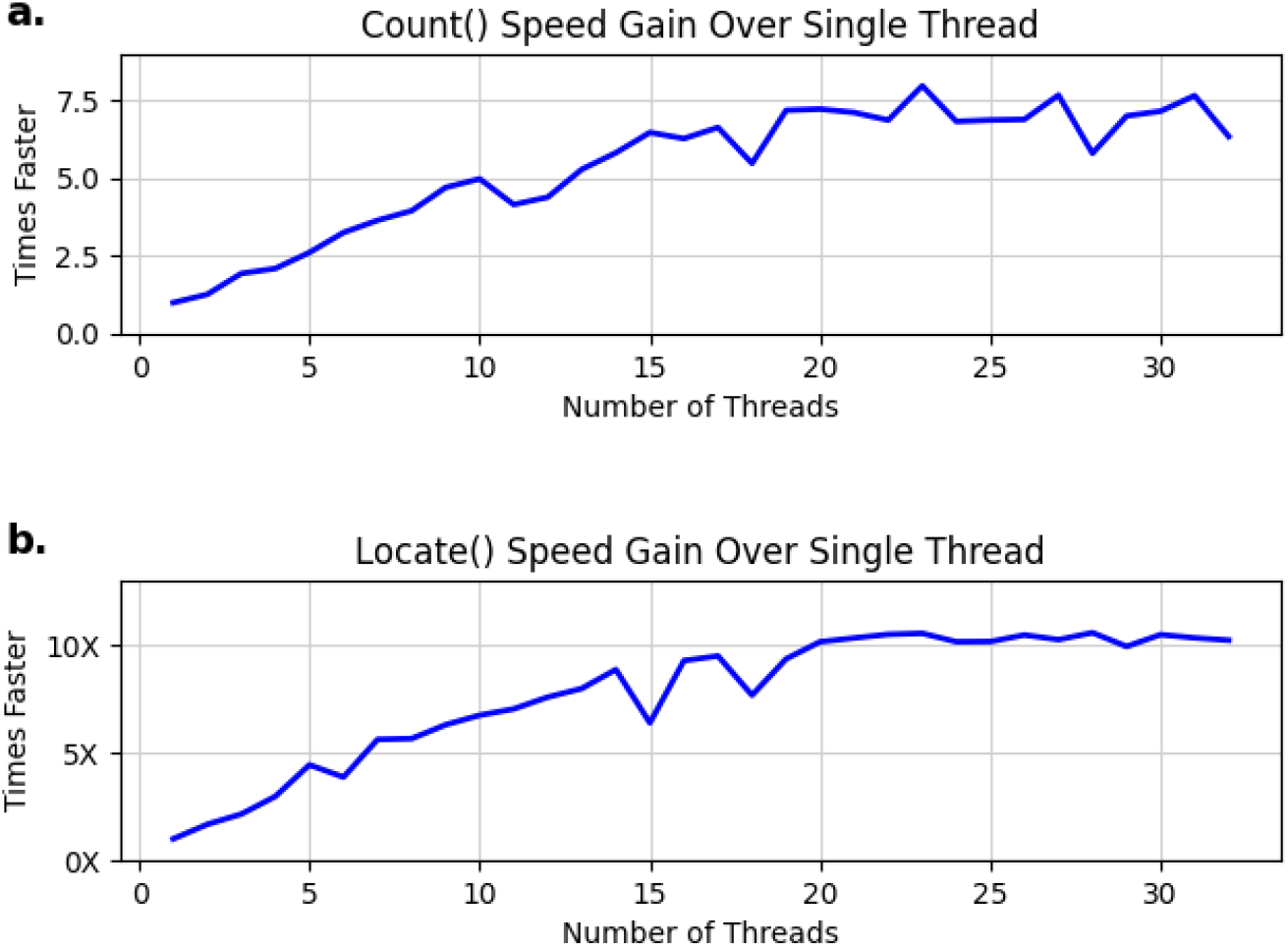
Parallel scaling. AWFM-index nucleotide Locate() search times for 1 million length 14 queries, parallelized with varying numbers of threads, from single-threaded search up to search using 42 threads against a target of 1 billion nucleotide-long target with a suffix array compression ratio of 4. (a) Increase in speed for the count() command, relative to single-threaded search. (b) Increase in speed for the locate() command, relative to single-threaded search.

### Prefetch Directives

The efficacy of data prefetch directives was analyzed by timing nucleotide locate() functions with each prefetch hint, and with prefetching directives disabled. Prefetch hint directives are used to tell the CPU which levels of cache to store the data in. All prefetching hints were shown to improve overall performance by a small amount, but non-temporal prefetching (MM HINT NTA) was shown to be fastest over multiple trials at a performance gain of 1.4%. Since the performance difference is minimal, we consider manual prefetching to not be a major contributor to AWFM-index’s overall performance.

## Discussion

We developed AWFM-index to be a lightweight, performant, easy-to-use library that simplifies the inclusion of fast pattern matching into bioinformatics software. Our implementation leverages a custom data layout and SIMD vectorized character comparison instructions to produce highly efficient symbol counting for nucleotide and amino acid alphabets. Combined with a pre-computed k-mer lookup table and out-of-the-box parallelism, the result is a library that provides very fast locate() and count() queries with very little development effort in the client. In addition to single-command search for full query sequences, the AWFM-index API also exposes stepwise iterative search functionality, so that clients can exert fine-grained control over FM index search steps, for example in support of back-tracking for inexact search as used in [2, 1].

AWFM-index offers good runtime performance relative to the mature SeqAn3 implementation, at the cost of an elevated memory footprint. Considering the now-ubiquitous availability of large memory systems, we expect that the runtime-memory tradeoff of the AWFM-index will be attractive to many developers. Even in low-memory systems, AWFM-index is still able to perform well using fully-sampled suffix arrays, particularly if the index resides on a low-latency solid-state drive.

While we expect AWFM-index to be immediately applicable in its current form, we note two potential improvements that will improve the future value of the library. The first of these is support for bi-directional FM-index search [24]. The bi-directional FM-index supports updates to the range of matching substrings by extending an existing substring [SP..EP] range with either a suffix or prefix symbol, and achieves this by supplementing the data structure with a single additional BWT over the reversed sequence T. Adding bi-directional search functionality to the library will improve it’s applicability to some special-case pattern matching applications such as [25].

The second improvement will extend the performance benefit of the k-mer lookup table. As described above, the BWT range of a query with the same length as the k-mer lookup table (e.g. nucleotide search for a length-12 query) is identified with a single memory access. Conversely, search for a slightly shorter k-mer (e.g. length 11 nucleotide query) does not use the k-mer lookup table, and thus receives no search shortcut (see Table 2, Fig 4). Future work on AWFM-index will enable application of the k-mer lookup table for queries shorter than k. For instance, consider the length-5 nucleotide query “CGTAG”, and a lookup table storing all length-7 nucleotide suffixes. Since all suffixes in the BWT are sorted, suffixes that begin with “CGTAG” will be found between (i) the start of the range for the k-mer extended with lowest rank non-sentinel symbol (here, “CGTAGAA”), inclusively, and (ii) the start of the range for the k-mer lexicographically one higher, extended with the lowest rank non-sentinel symbol (here, “CGTA<u>T</u>AA”), exclusively. However, use of these longer strings as proxies during the identification of the BWT range fails to account for the possibility of a sentinel symbol, which may introduce a non-matching string into the proxy range. Since a BWT is guaranteed to only contain a single sentinel symbol at the end of the sequence, the last few symbols of the original text T can be kept along with the k-mer lookup table, and used to remove this matches from a range list. A more thorny problem arises when the short query k-mer ends with a symbol of the highest rank, non-ambiguity symbol (nucleotide T or amino acid Y), as the lookup table does not have a higher-rank symbol to use in selecting the top end of the range. One way to resolve this is to store all ranges in the k-mer table, including those that contain ambiguity symbols; however, including the ambiguity symbol X increases the table size appreciably, e.g. a table of all length-12 nucleotide k-mers takes 16 · 4^12^ = 268MB, while the same table that also stores ambiguity characters takes 16 · 5^12^ = 3.9GB. Other possible solutions to this issue include using the partial k-mer lookup table for all queries that don’t end with a nucleotide T or amino acid Y as described above, and querying using the traditional backwards search for those queries that do. Perhaps the simplest solution to this problem involves keeping multiple k-mer lookup tables of varying k-mer lengths. Since the memory footprint of the table grows exponentially with the length of the k-mer, a table made from shorter k-mers will use much less memory: while an index containing a length-12 k-mer table takes 268MB, adding a length-10 and a length-6 table would cumulatively add only (16 · 4^10^) + (16 · 4^6^) = 16.8MB of memory, but would improve runtime for any queries length 6 to 11. We plan to update AWFM-index to support bi-directional indexes and using k-mer lookup tables for small queries in a future library release.

## Availability and requirements

**Project name:** AWFM-index library

**Project home page:** https://github.com/TravisWheelerLab/AvxWindowFmIndex

**Operating system(s):** Unix/Linux

**Programming language:** C

**Other requirements:** None

**License:** BSD-3-Clause

**Any restrictions to use by non-academics:** None

### Ethics approval and consent to participate

Not applicable

### Consent for publication

Not applicable

### Availability of data and materials

Data used to produce figures for this manuscript can be found at http://wheelerlab.org/publications/2021-AWFM-Anderson/Anderson_suppl.tar.gz.

### Funding

This work was supported by NIH grant P20 GM103546 (NIGMS) and DOE grant DE-SC0021216.

### Author’s contributions

TA devised all algorithms and code optimizations, implemented all code, performed all benchmarking experiments, and led development of the manuscript. TJW introduced the problem to TA, provided guidance on use cases and experimental design, and assisted with writing the manuscript.

## Acknowledgements

We thank Robert Hubley for beta testing AWFM-index and suggesting improvements to the library’s build process, as well as George Lesica for his help in improving the library API and implementing Cmake support. We also gratefully acknowledge the computational resources provided by the University of Montana’s Griz Shared Computing Cluster (GSCC), and thank the reviewers for helpful suggestions that have improved the quality of this manuscript.

